# Metabolic flexibility among fiber degraders supports gut microbiome stability across host diets in the American cockroach

**DOI:** 10.1101/2024.10.31.621369

**Authors:** Amber N. DePoy, Helen E. Wall, Kara A. Tinker, Ian M. Black, Elizabeth A. Ottesen

## Abstract

Scavenging omnivores face challenges in maintaining gut microbiome homeostasis due to a highly variable diet. Work in mammalian models has highlighted the importance of fiber-degrading taxa and their utilization of dietary polysaccharides and host glycans in structuring gut microbiome composition and function. However, less is known about biological mechanisms supporting gut microbiome homeostasis in insects. Here, we present a metatranscriptomic analysis of hindgut microbial transcriptional responses to diverse host diets in the American cockroach *(Periplaneta americana*), a hindgut-fermenting and omnivorous insect. We analyzed microbial community transcriptomes from hindguts of cockroaches fed high-fiber, high-protein, high-fat, and starvation diets. Microbial taxa predicted to be primary fiber degraders increased transcription of genes associated with utilization of dietary polysaccharides in insects fed a high-fiber diet. In contrast, these taxa responded to starvation and low fiber diets with upregulation of transcripts associated with degradation of host glycans. Analysis of N- and O-glycans in the cockroach midgut confirm the presence of predicted substrates for these pathways. As a result, we hypothesize that peritrophic matrix glycans serve as a critical carbohydrate subsidy supporting insect gut microbial metabolism in the absence of dietary fiber. Observed stability in gut microbiome composition and the transcriptional activity of many secondary fermenters and hydrogenotrophic taxa in the gut suggests that this strategy can be highly successful. Together, this work supports a novel physiological role for the peritrophic matrix of insects and other invertebrates in supporting gut microbiome homeostasis across dietary shifts.

## Introduction

The gut microbiome shares an intimate relationship with its host, impacting digestion, nutrition, immunity, and more [1–5]. The host can modify the gut microbiome through various pathways [6–10], but diet is considered one of the most direct drivers of gut microbiome composition [11–15]. Much of the work studying the effect of diet on the gut microbiome has been done in mammalian hosts, such as mice [16–18], humans [11,19–21], and other mammals [22–24]. These mammalian studies have shown that dietary fiber and plant polysaccharides play a particularly important role shaping gut microbiome composition [25–27]. Despite the importance of diet in shaping the gut microbiome, multiple long-term studies of individual humans have shown stability over time [20,28–30]. Short-term dietary interventions in humans often fail to elicit shifts in the gut microbiome composition more than pre-existing individual-to-individual differences [28,31,32]. These results emphasize the need for an understanding of microbe-microbe or host-microbe interactions that may assist in the maintenance of gut microbiome homeostasis.

The gut microbiome of many insects is less diverse and more variable than that of mammals [2], however there are numerous exceptions, including dung beetles, termites, ants, and cockroaches [33–38]. Insect models such as these provide an opportunity to explore diverse and evolutionarily ancient biological mechanisms supporting host-gut microbe symbioses. In this work, we explore gut microbiome homeostasis in response to host dietary shifts in the omnivorous American cockroach (*Periplaneta americana*). The American cockroach hindgut microbial community is highly diverse and includes many taxa from clades found in mammalian and human guts [33,39–41]. These taxa include those in the abundant Bacteroidota and Bacillota phyla that have similar taxonomic and functional profiles to their sister clades found in mammalian guts [42]. Multiple studies have demonstrated that, like humans, the cockroach gut microbiome exhibits stability across highly divergent diets [33,40,43,44]. This stability is likely advantageous to their lifestyle as scavenging omnivores, but host and microbial activities supporting this stability are poorly understood.

Our goal of this study was to explore the microbial activities underlying compositional stability with host diet changes, with a particular focus on understanding metabolic shifts supporting the ability of key gut taxa to persist in the absence of a dietary source with preferred growth substrates. We examined hindgut microbial community activity using metagenomic and metatranscriptomic approaches in cockroaches fed a high-fiber, high-protein, high-fat, and starvation diet. Metagenomic analyses confirmed taxonomic stability, while metatranscriptomic profiles showed plasticity in transcriptional activity among key taxa, particularly primary fiber degraders. Analysis of carbohydrate active enzyme expression highlights a role for host glycans in supporting fiber degraders in low fiber diets. Together, these results provide new insight into microbial responses to dietary shifts that support gut microbiome homeostasis.

## Materials and Methods

### Diet manipulation and sample collection

Cockroaches (*Periplaneta americana*) are maintained in mixed age, mixed sex colonies in aquarium tanks at room temperature on a diet of dog food *ad libitum*. Each tank is provided with corn cob bedding, cardboard tubes for nesting, and a cellulose sponge saturated with water. For experiments using controlled diets, each treatment group was housed in a single plastic tank that contained pebbles for bedding, a large weigh boat for shelter, and food and water in weigh boats. Food, water, and weigh boats were changed as needed, and any ootheca or deceased cockroaches were removed daily. Treatments included bran (Bob’s Red Mill Organic High Fiber Oat Bran Hot Cereal), butter (Kroger Unsalted Butter Sticks), tuna (StarKist Selects Low Sodium Chunk Light Tuna in Water), and starvation (Table 1).

**Table 1.** Nutrient information for 100g serving of diet treatment. All cockroaches were raised on our laboratory control diet of dog food, composed of 21% protein, 8% fat, and 6% fiber.

| Diet Treatment | Calories | Protein (g) | Carbohydrate (g) | Fat (g) | Fiber (g) |
| --- | --- | --- | --- | --- | --- |
| Bran (high-fiber) | 375 | 17.5 | 67.5 | 5 | 17.5 |
| Butter (high-fat) | 714 | 0 | 0 | 79 | 0 |
| Tuna (high-protein) | 107 | 27 | 0 | 1 | 0 |
| Starvation (starve) | N/A | N/A | N/A | N/A | N/A |

After 14 days on the assigned diet, all cockroaches were sacrificed and their hindgut microbiota preserved for DNA or RNA extraction. Individual cockroaches were removed from tanks, weighed, and placed on ice in sterile culture plates. Once sufficiently torpid, cockroaches were dissected, and the entire gut was removed. Any visible debris, including fat bodies or exoskeleton, was removed with forceps. The hindgut was then separated from the rest of the gut using a scalpel and submerged in either 100 μL of 1XTE buffer or RNALater (Ambion, Austin, TX, USA). Hindgut samples placed in 1XTE buffer were immediately stored at −80°C. Hindgut samples placed in RNALater were crushed with a sterile glass stirring rod. The suspended gut lumen was then aspirated and stored at −80°C.

### DNA and RNA extraction, sequencing library preparation

Microbial DNA and RNA were extracted from hindgut samples as outlined previously in [42]. Briefly, DNA and RNA were extracted using a modified version of the EZNA Bacteria Kit and HP Total RNA Kit (Omega Biotek, Norcross, GA), respectively. For metatranscriptome sequencing, RNA libraries were prepared using the NEBNext Ultra Directional RNA Library Prep Kit for Illumina (NEB) following product guidelines with an RNA fragmentation time of 10 min. For metagenome sequencing, DNA libraries were prepared using NEBNext Ultra II DNA Library Prep Kit for Illumina (New England BioLabs Inc) and dual index primers from the NEBNext Multiplex Oligos for Illumina (New England BioLabs Inc.). Size selection was done using a protocol adapted from [45]. The prepared RNA library was sequenced (Illumina HiSeq) by the Georgia Genomics and Bioinformatics Core (Athens, GA), while the metagenome pool was sequenced (Illumina NovSeq) by Novogene Corporation Inc. (Sacramento, CA).

Raw sequences were checked for low-quality regions using FastQC and MultiQC. Adapter sequences were trimmed using BBDuk from BBTools. Ribosomal RNA sequences were removed using SortMeRNA [46]. *Periplaneta americana* [47] and *Blattabacterium* [48] sequences were removed using BBSplit. Paired-end reads were merged using BBMerge and reads that did not successfully merge were rescued by pseudo-merging the forward read with an internal spacer of 10 Ns and the reverse complement of the reverse read. This approach allowed the processing of pseudo-merged reads through downstream scripts in the same manner as merged reads.

### Construction of reference genome clusters for key taxonomic clusters

An obstacle to RNA-seq analysis of gut microbiota is the relative lack of available reference genomes. To surmount this problem, we utilized a pangenome mapping approach based loosely on that employed in [49]. This approach uses available reference genomes associated with taxonomic clusters of interest to construct genome clusters for analysis of group-level transcriptional activity. Reads were mapped to pangenomes constructed from clusters of closely related genomes that show high levels of co-occurrence in the metagenome and metatranscriptomes (e.g., Figures S1-S2). These analyses allowed us to compile genetic information from multiple closely related reference genomes while minimizing ambiguity of read mapping in conserved regions. Pangenomes were constructed for 18 genome clusters (see Table S1 for a full list of reference genomes used for constructing each cluster). Genome sequences for inclusion in each genome cluster were identified using a variety of metrics, including taxonomic annotation, phylogenetic information from 16S rRNA gene and genome trees, abundance in metagenome and metatranscriptome based on RefSeq homology searches as described below, and co-correlation analyses.

Pangenomes of orthologous gene clusters for each taxon were built with Anvi’o [50]. We used *anvi-pan-genome* with DIAMOND [51] to calculate genome similarity within the taxon and bin individual genes into gene clusters using the MCL algorithm [52,53]. Functional annotations for gene clusters were found using DIAMOND [51] to match functions and pathways in KEGG [54], COG [55,56], and CAZy [57]. Additional annotations from a DIAMOND search of each taxon’s reference genomes to the Transporter Classification Database (TCDB) were added to gene clusters [58].

Metatranscriptome and metagenome reads were translated and mapped to protein sequences in RefSeq Prokaryotic Genomes (accessed 11/2021) [59] and to our dataset of 96 single-cell genomes from cockroach gut microbes [42,60] using DIAMOND [51]. The top-scoring hit (based on bit score) and all equal hits were kept. Custom scripts were used to map reads to genome clusters and build a per-sample gene cluster hit count table. See supplemental Table S1-S3 for read tracking through the analysis steps.

### Analysis of genome cluster abundance and transcriptional activity

To calculate genome cluster abundance, the number of metatranscriptome and metagenome read hits to each taxon was divided by the number of reads in that sample with hits to the combined RefSeq and SAG reference database. Weighted Bray-Curtis dissimilarities were calculated in R on whole-community taxonomic tables using *vegdist()* from the vegan package [61]. Differences in dissimilarity values between diets were tested for significance in R with the Kruskal-Wallis method followed by Dunn’s test with Bonferroni correction (*dunnTest()* from FSA package and *kruskal.test()* from stats package, respectively). Permutational multivariate analysis of variance (PERMANOVA) was carried out on dissimilarities using *adonis2()* from the vegan R package with host diet as the factor of variation. Redundancy analyses (RDA) were done with variance stabilized data normalized with DESeq2 [62,63]. For all taxa, DESeq2 [62,63] was used for pairwise comparisons between diet treatments to find significantly differentially expressed gene clusters between diets.

## Results

### Taxonomic profiles and overall community composition

To address gut microbiome stability and the functions underlying that stability, we sequenced metagenomes and metatranscriptomes from cockroaches fed four distinct dietary conditions: bran (high-fiber), tuna (high-protein), butter (high-fat), and starvation. Metagenomic reads were assigned to a diverse set of families, including Bacteroidaceae, Dysgonomonadaceae, Tannerellaceae, Lachnospiraceae, and Desulfovibrionaceae (Figure 1). Relative abundances of these families changed very little between diet treatments. Alpha diversity (Shannon index) did not vary significantly across the diets (*p* = 0.43) (Figure S3). PERMANOVA analysis of beta diversity using weighted Bray-Curtis dissimilarities based on taxonomic profiles confirmed that diet was not significantly correlated with overall metagenome community composition (r^2^ = 0.18, *p* = 0.24) (Figure 1), although we did observe significant enrichment/depletion of individual taxa (Figure S4). Much of this taxa enrichment/depletion appears to be associated with increased intra-treatment variability and associated ‘blooms’ of multiple taxa in individual samples within the butter diet (Figure S1, Figure S5).

**Figure 1.**
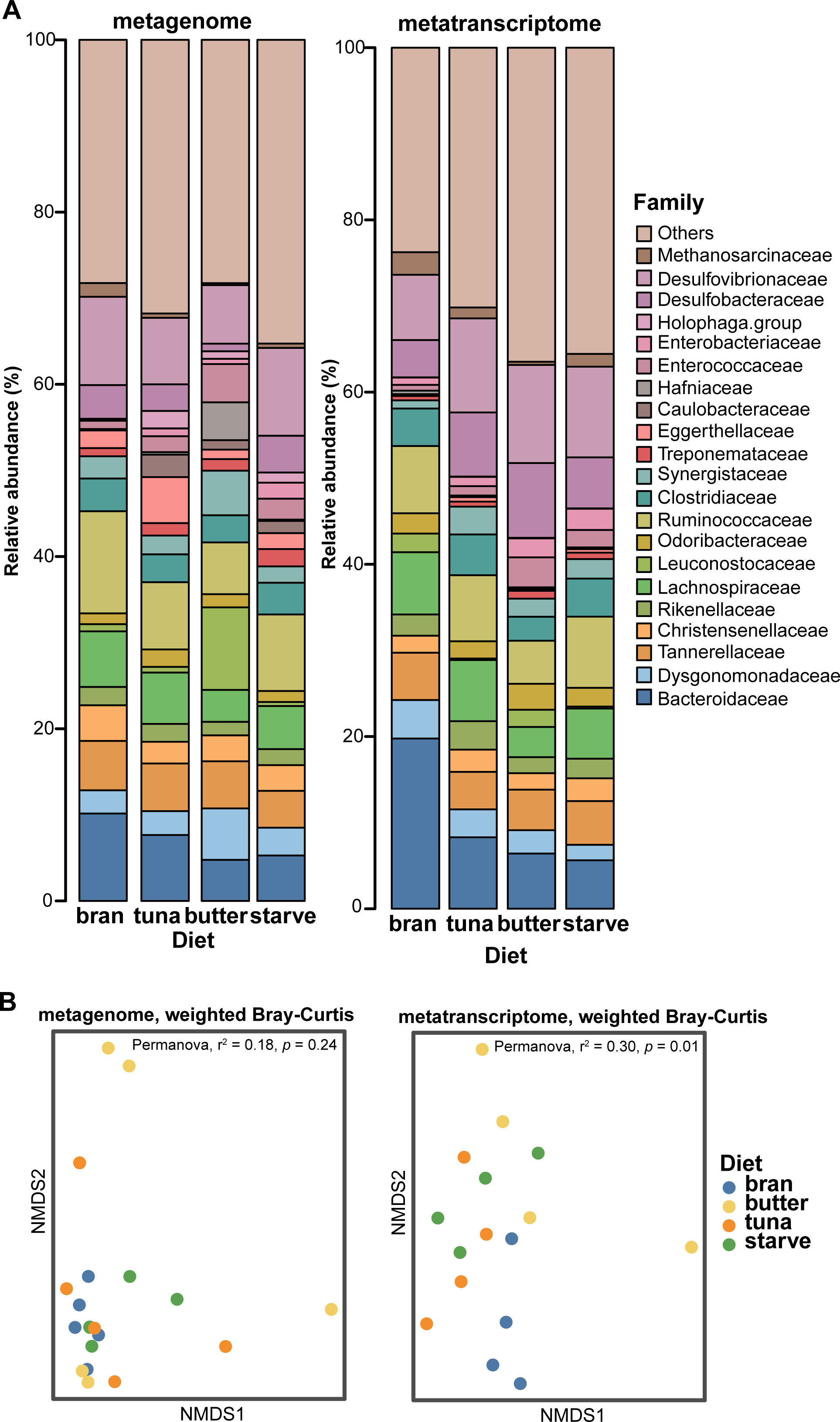
Taxonomic composition of metagenomes and metatranscriptomes. A) Bar plot of relative abundance (%) of families in the metagenome and metatranscriptome datasets. Each bar represents an average relative abundance of 5 replicates (metagenome) or 4 replicates (metatranscriptome) per diet. Families with a maximum relative abundance of at least 5% in any sample of either the metatranscriptome or metagenome dataset are included in the taxonomy. B) Non-metric multidimensional scaling (NMDS) using a weighted Bray-Curtis dissimilarity metric for the metagenome and metatranscriptome datasets, labeled accordingly.

In contrast, diet was a significant driver of metatranscriptome composition (r^2^ = 0.30, *p* = 0.01) and alpha diversity (Shannon index) (*p* = 0.005; bran vs. starve, *p* = 0.02) (Figure 1, Figure S3). DESeq2 analysis of taxonomic composition showed significant enrichment of diverse Bacteroidaceae in the bran diet compared to the other diets, as well as enrichment of Lachnospiraceae, Leuconostocaceae, and Clostridiaceae (Figure S4). Few taxa were enriched in the other diets (Figure S4).

### Response of individual taxa to host diet treatment

Transcriptional profiles of each taxon were analyzed using redundancy analysis (RDA), which demonstrated that diet was significantly associated with overall transcriptional activity in most taxa, excluding Bacteroidales, Enterococcaceae, Lactobacillaceae, and *Desulfosarcina* (Figure S6). When ‘diet’ was modeled as a factor, it explained 8.3% (Bacteroidales) to 21.9% (Lactobacillaceae) of the variance (Figure S6). For many organisms, the presence/absence of fiber alone (bran vs. all other diets) explained a comparable amount of transcriptional variance to ‘diet’ as a factor. This emphasizes the effect and importance of dietary fiber in shaping microbial transcriptional profiles.

To characterize microbial responses and the extent of responses to host diet changes, we used DESeq2 to identify genes that displayed significantly altered expression in pairwise comparisons of the diet treatments. A combined total of 2,684 gene clusters were changed significantly between host diets (Figure 2). Members of the Bacteroidota phylum altered expression of the greatest number of genes, with *Bacteroides*, *Dysgonomonas,* and *Parabacteroides* having far more than every other group (Figure 2, Figure S7, Table S4). Many genes significantly changed expression between the high-fiber (bran) diet compared to each of the other diets (Figure 2). Two Bacillota taxa, Oscillospiraceae_2 and Lactobacillaceae, had high numbers of significantly responding genes (Figure S7, Table S5). However, other Bacillota taxa had fewer differentially transcribed genes. Chemolithotrophic groups (sulfate reducers and methanogens) had a range of significantly changing genes but showed less of a response with gene expression changes than other taxa (Figure S7, Table S6).

**Figure 2.**
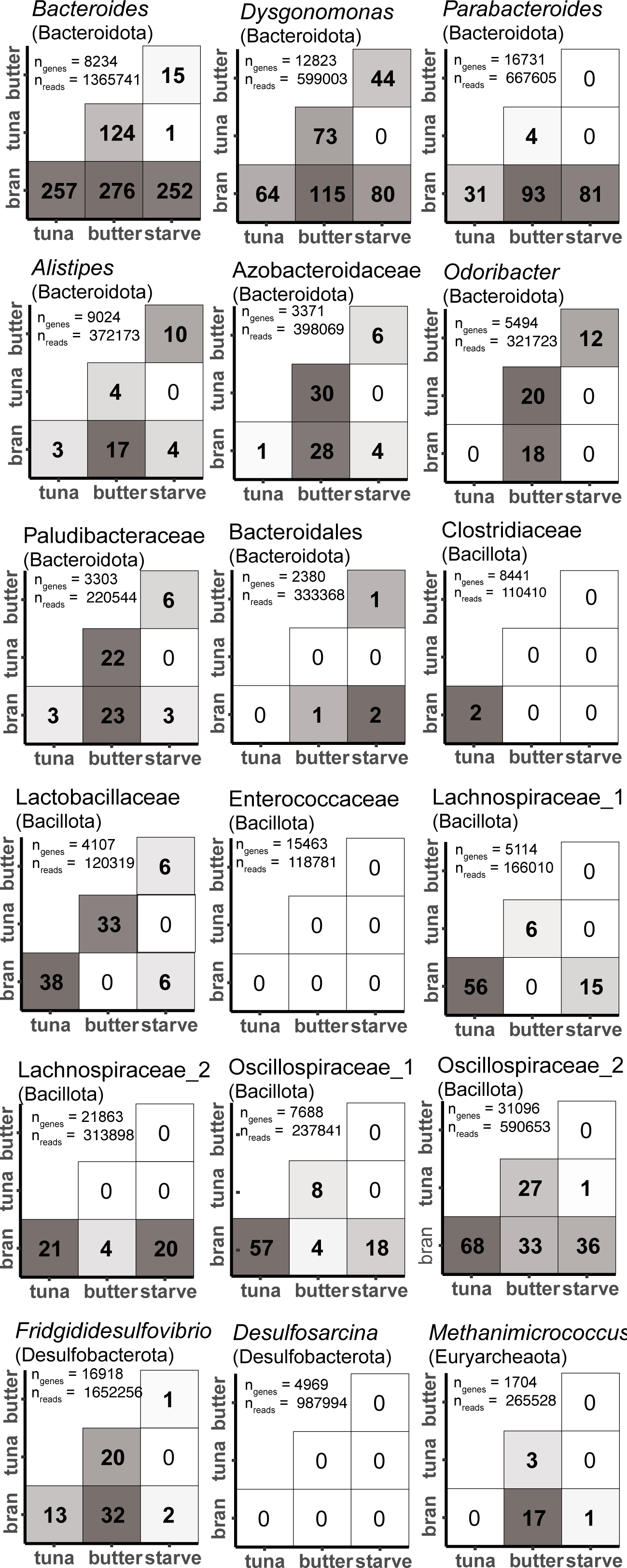
Number of differentially expressed genes identified by DESeq2. Each number within each square is the number of differentially expressed genes when those two diets were compared.

### Carbohydrate active enzymes (CAZyme) utilization between the host diets

Given the importance of fiber in shaping transcription, we analyzed genes active in the breakdown of dietary polysaccharides (CAZymes). CAZyme gene expression (% of all mapped transcripts) was highest in the bran diet (2.1% of all transcription in bran) and decreased with the amount of fiber present in the diet (Figure 3B). Despite starvation having no dietary fiber input, CAZyme gene expression remained high under starvation, still accounting for 0.99% of all transcription (Figure 3B). *Bacteroides*, *Dysgonomonas*, and *Parabacteroides* produced the highest number of CAZyme transcripts, emphasizing their role as primary degraders (Figure 3).

**Figure 3.**
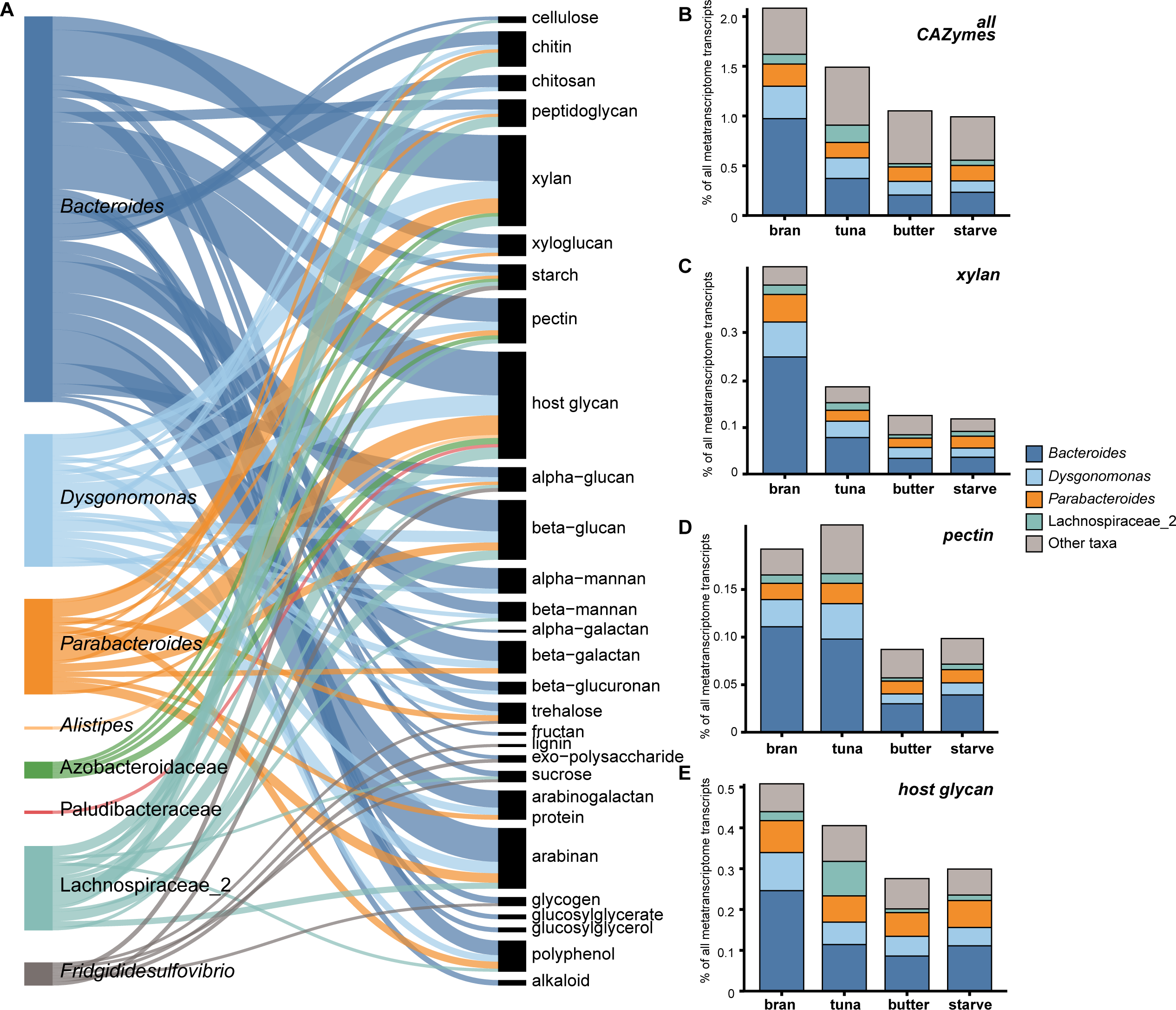
CAZyme gene expression across the gut microbiome. A) Sankey diagram of the summed relative abundance of CAZyme genes (> 0.01% relative abundance) associated with the breakdown of substrates. Relative abundance was calculated as a proportion of all transcripts in each sample to allow for comparisons between taxa. Lines of the Sankey diagram represent the summed relative abundance of the average across all diets, while the stacked bar plots represent summed relative abundance of transcripts in each fiber substrate averaged for each diet. B) Bar plot of transcription of B) all, C) xylan, D) pectin, and E) host glycan CAZymes across all diets. Note that different scales were used on the y-axes of the bar plots to account for differences in expression levels of each of the substrate CAZyme families.

We used dbCAN-seq [64] to associate CAZyme family annotations with potential substrates (Figure 3). These annotations can be somewhat ambiguous as fiber substrate breakdown can overlap between CAZyme family classifications, but it remains a valuable way to identify changes in putative substrate profiles across diets. CAZymes targeting three major substrate types were expressed at high levels: xylan, pectin, and host glycans (Figure 3A). Xylan-targeting CAZymes were expressed 2.4-fold more in the bran diet compared to the other diets (Figure 3C). Pectin breakdown genes were expressed 2-fold more in tuna diet than butter and starve but only slightly more than in bran (Figure 3D). Host glycan-targeting CAZymes were among the most highly expressed across all conditions (Figure 3E). While all CAZyme groups showed decreased expression in starvation (and butter), host glycan targeting CAZymes showed smaller diet-dependent fluctuations than other CAZyme groups.

### Patterns of CAZyme and carbohydrate utilization gene expression in Bacteroides

As one of the most active genome clusters in the metatranscriptome and the largest source of CAZyme transcripts, we performed a more in-depth analysis of CAZyme expression and carbohydrate utilization gene expression in *Bacteroides*. For these analyses, we expressed transcription of each fiber substrate metabolism genes as a fraction of *Bacteroides* transcripts. This standardization method allows us to examine changes in transcriptional activity within *Bacteroides* (equivalent to single-organism RNA-seq) in isolation from changes in the relative abundance of this organism within the metatranscriptome.

For *Bacteroides*, overall transcription of xylan targeting CAZymes was highest in the bran diet (Figure 4A). Several CAZymes that target the xylan backbone (GH10 and GH3) were significantly upregulated in bran (Figure 4A), as were CAZyme families targeting arabinose (GH51) and glucuronate (GH115) side chains of xylan. Although not significant, CE1 and GH67 (required for removal of acetyl groups and glucuronic acid side chains) also had greater transcription in the bran diet (Figure 4A). While xylan gene expression remains high in starvation, much of this signal is associated with transcription of broad specificity CAZyme families (i.e., GH2, GH3) that are associated with utilization of a wide range of substrates. We also examined KEGG orthologs associated with processing of sugars released during xylan breakdown, focusing on early processing steps prior to entry into shared metabolic pathways. *Bacteroides* expressed two genes for xylose processing into xylulose-5P (for entry into the pentose phosphate pathway) at high levels the bran diet (Figure 4D). *Bacteroides* also expressed L-arabinose isomerase (K01804) at high levels in both bran and tuna diets (Figure 4G). Arabinose can be released from either arabinoxylan or pectin processing.

**Figure 4.**
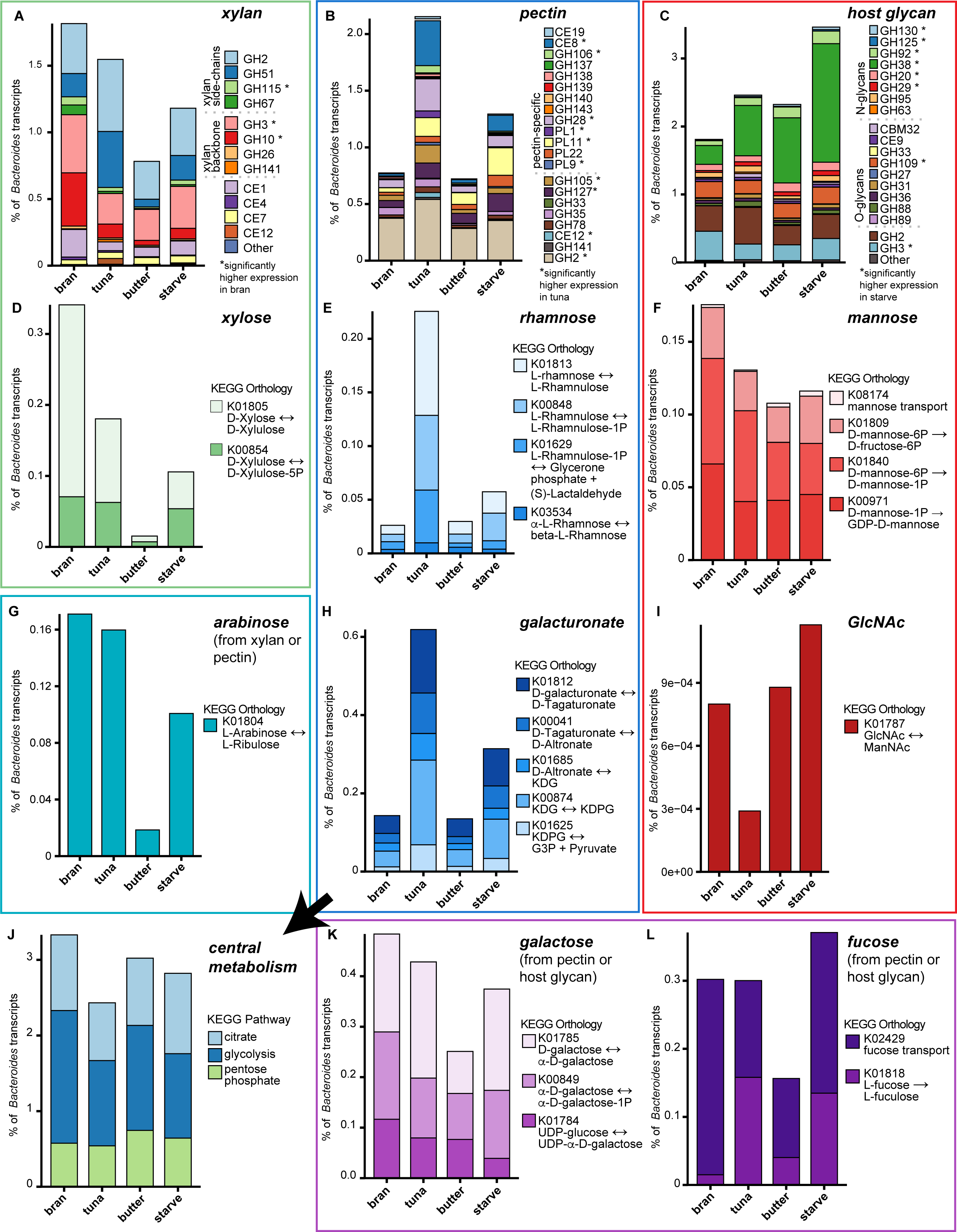
Xylan (A), pectin (B), and host glycan (C) utilization by *Bacteroides*. Bar plots of average (n = 4) relative abundance (%) of CAZyme families and KEGG orthologs associated with polysaccharide degradation. Each bar plot is labelled with its associated substrate of degradation: xylose (D), rhamnose (E), mannose (F), arabinose (G), galacturonate (H), GlcNAc (I), galactose (K), and fucose (L). Note each bar plot has a different y-axis scale to emphasize the difference between expression levels between diets. Monosaccharide degradation following its respective polysaccharide are together in a box, while overlapping monosaccharides are separately boxed and colored. Abbreviations, galacturonate: KDG = 2-Dehydro-3-deoxy-D-gluconate, KDPG = 2-Dehydro-3-deoxy-6-phospho-D-glyconate, G3P = D-Glyceraldehyde 3-phosphate. All monosaccharide degradation pathways feed into central metabolism pathways (J) shown in lower left bar plot.

*Bacteroides* responded to the tuna diet with increased expression of pectin targeting CAZymes (Figure 4B). There were ten CAZyme families that were significantly upregulated in tuna, six of which were specific for pectin (Figure 4B), including CE8 demethylases, GH105 and GH106 for depolymerization of the pectin backbone, and polysaccharide lyases PL1, PL11, and PL9 (Figure 4B). Although not specifically associated as a pectin CAZyme family, GH51 (L-arabinfuranosidase) was also significantly upregulated in the tuna diet (Figure 4A). After initial pectin breakdown, the main monosaccharides released for further metabolism include rhamnose, galactose, galacturonic acid, fucose, and arabinose. *Bacteroides* expressed rhamnose and galacturonic acid metabolism genes more in the tuna diet compared to the bran diet (8.6 and 4.3-fold more, respectively) (Figure 4E, 4H). Galactose metabolism genes did not show the same increase in the tuna diet but were highly expressed in most diets (Figure 4K).

Relative to total organismal transcriptional activity, *Bacteroides* showed 1.91-fold higher expression of host glycan degradation CAZymes in starvation than in the bran diet (Figure 4C). Eight CAZyme families associated with host glycan utilization were significantly upregulated in starvation vs. bran. Most of these CAZymes were associated with N-glycan degradation (Figure 4C). *Bacteroides* increased expression of GH38 (α-mannosidases) 6.3-fold more in host starvation compared to bran, suggesting target of mannoses (Figure 4C). In addition to GH38, other CAZyme families with mannosidase activity (GH130, GH125, GH92) were significantly upregulated in starvation compared to bran. *Bacteroides* also targeted fucose with two CAZymes, one of which (GH29) was significantly upregulated in starvation. CAZyme families targeting O-glycans were expressed to a lesser extent. Cleavage of terminal α-GalNAc residues from O-glycans by GH109 was significantly upregulated in starvation. Monosaccharides from N-glycan degradation include mannose, fucose, and GlcNAc. Genes associated with mannose breakdown were transcribed under all conditions, although they were not significantly upregulated under starvation (Figure 4F). Although not significant, fucose metabolism genes were transcribed more in starvation and less in the butter diet (Figure 4L).

### Taxa showing functional stability among the host diets

While primary degraders showed changes in CAZyme gene expression and downstream utilization of the monosaccharides to each of the diets, these significant changes underscore the extent to which other taxa were less affected by host diet changes. Secondary degraders, including many organisms within the Bacillota, showed fewer significant differences in gene expression across diets, and the presence/absence of dietary fiber explained a smaller fraction of their overall transcriptional profile (Figure 2, Figure S6-S7). While the major fiber sources in each diet require unique carbohydrate active enzymes to degrade, this degradation produces a limited set of monosaccharides, which are ultimately metabolized via conserved core metabolic pathways. Summed gene expression among core metabolic pathways, such as glycolysis, pentose phosphate, and citrate pathways showed high and stable expression across the diets in most organisms (Figure S8).

Other groups that are not directly involved in polysaccharide degradation, including sulfate reducers (*Desulfosarcina*, *Frigididesulfovibrio*) and methanogens (*Methanimicrococcus*), expressed very few genes differentially between the diets (Figure 2). Despite an overall transcriptional abundance like that of key fiber degraders, *Desulfosarcina* did not differentially express any genes across the diets, while *Frigididesulfovibrio* differentially expressed only 60 genes. None of the significantly enriched genes for *Frigididesulfovibrio* were related to sulfate reduction. Of the differentially expressed genes, most were enriched in the tuna diet (28 out of 60 genes) compared to each of the other diets, including three genes associated with the taurine metabolism pathway (K03851, K03852). *Methanimicrococcus* differentially expressed few genes, with most gene expression changes identified in the bran vs. butter comparison (17 genes) (Figure 2). Ten of these genes were transcribed more in the bran diet including four hypothetical proteins, two coenzyme F420 hydrogenase subunits (frhA and frhG), cobaltochelatase subunit CobN, oxidoreductase hydrogenase small subunit, and respiratory nitrate reductase subunit gamma. Conversely, there were seven genes that were transcribed more in the butter diet compared to bran, including methanol:5-hydroxybenzimidazolylcobamide Co-methyltransferase, monomethylamine corrinoid protein, ethanolamine permease, acetate uptake transporter, and peroxiredoxin (Table S6).

### High-fat diet induces community stress response

Second to the high fiber diet, many of cockroach gut bacteria exhibited strong transcriptional responses to the butter (high-fat) diet, including some taxa that showed minimal responses to other dietary treatments (Figure 2). Across all taxa, there were 1130 genes that were significantly differentially expressed between butter and other diets. Of these genes, 38% (410 genes) were enriched in the butter diet compared to the other diets. Many of these genes were annotated as chaperones and folding catalysts including DnaJK chaperones, heat-shock proteins, peroxiredoxin, and thioredoxin (Figure S9). 11 out of 18 taxa exhibited significantly increased expression in at least one of these transcripts, and 6 of the remaining taxa showed increases below the significance threshold.

## Discussion

In this study, we examined cockroach gut microbiome transcriptional activity under diverse host diets to better understand metabolic responses to changing dietary inputs and how they may support gut microbiome homeostasis. Our previous work showed that the American cockroach gut microbiome demonstrates high taxonomic stability following dietary shifts [33], even following such extreme treatments as starvation for up to 60 days. Much of the stability observed in the American cockroach bears similarities to the long-term taxonomic stability seen in humans [20,28–30]. However, factors contributing to this stability are poorly understood.

Metagenomic analysis supported 16S rRNA-based findings that the cockroach gut microbiome showed high taxonomic stability across diets (Figure 1). However, we observed diet-driven changes in both the relative abundance of different taxonomic groups in the metatranscriptome and the transcriptional profiles of individual organisms. *Bacteroides*, *Dysgonomonas*, and *Parabacteroides* have the greatest plasticity in gene expression across host diets, suggesting that they play an important role in shaping gut microbiome responses to host diet (Figure 2). These taxa have previously been hypothesized to be primary fiber degraders based on the number of CAZymes (carbohydrate active enzymes) present in their genomes and the level at which these CAZymes are transcribed (Figure 3) [42].

Analysis of transcripts annotated as CAZymes showed diet-associated differences in polysaccharide utilization profiles. The oat bran diet had the most dietary fiber content (Table 1) and contains arabinoxylan along with beta-glucans [65]. We found that this bran diet led to high expression of CAZyme families associated with arabinoxylan utilization. Highly transcribed CAZymes including enzymes associated with side chain removal and degradation of main chain xylose-xylose bonds, including GH2, GH3, GH10 and CE1. Several *Bacteroides* isolates from the human gut are also known to highly express these xylanase CAZyme families when grown on xylan [66–68].

The tuna diet was associated with unexpectedly high transcription of CAZymes associated with pectin utilization, including CE8 (pectin methyl-esterase), which has been shown to be particularly important in pectin metabolism by *Bacteroides thetaiotaomicron* [69]. We note that the canned tuna used includes vegetable broth as a flavor enhancer and hypothesize that gut bacteria may be responding to pectin present in this additive. In addition, there have been reports of rhamnose, xylose, mannose, glucuronic acid, and galacturonic acid in processing Skipjack tuna by-products, which suggests that glycans naturally present in tuna may be inducing expression of CAZymes typically associated with pectin degradation [70].

The butter diet was associated with a decrease of CAZyme transcripts (Figure 3) as well as widespread transcriptional upregulation of stress response genes including RND efflux pumps, oxidative stress, and heat shock proteins (Figure S9). Two notable proteins that showed increased expression in the butter diet, GroEL and dnaK, are known to play an important role in bacterial survival during chemical, heat, or oxidative stress [71–73]. While additional work is needed to determine if high fat diets induce host oxidative stress in the cockroach, high-fat diets in mouse models have been associated with oxidative stress [74] and changes in redox state in gut epithelial cells [75,76]. The butter diet in the cockroach also appears to reduce gut microbiome stability, with blooms of facultative anaerobes such as *Enterococcaceae* and *Lactobacillaceae* (*Leuconostocaceae*) observed in metagenomic analyses of a subset of cockroaches eating butter (Figure S4-S5). These results are notably similar to studies that have found blooms of organisms [18,77,78] and growth of facultative anaerobes, including *Enterococcaceae* and *Lactobacillaceae* spp. in mice eating high-fat diets [79–81]. Studies in humans have also indicated greater variability in community profiles of individuals eating high-fat diets [82]. Together, these results are consistent with a pattern of high fat diet causing oxidative stress in the gut as well as blooms of oxygen-tolerant microbiota and a reduction in the abundance and activity of fiber degrading bacteria. The remarkable similarity in responses to high fat diet between cockroaches and mammalian models suggests that the cockroach gut may represent a useful model for exploring these linkages further.

A major goal of this work was to identify microbial metabolic activities that support the observed persistence of diverse gut microbes under starvation conditions as well as other diets lacking preferred growth substrates. A key finding was that overall CAZyme transcription remained relatively high during host starvation, including strong and stable transcription of CAZymes for utilizing host glycans (Figure 3). This signal was particularly notable in *Bacteroides,* with eight CAZyme families associated with host glycan utilization showing significantly higher expression under starvation compared to bran and/or butter (Figure 4C). The utilization of host glycans has been widely reported in the gut symbionts of mammals such as humans and mice, which helped identify host mucins as an important substrate for gut fiber degraders in the absence of microbially accessible polysaccharides [83–85]. More broadly, host mucins have been shown to play a role in shaping symbiotic microbiota across the animal kingdom, from invertebrates such as corals, squid light organs, and insects to higher animals including zebra fish, mice, and humans [86–90]. However, it was not immediately clear that this strategy would be applicable to the cockroach hindgut microbiome, as the insect hindgut is surrounded by a chitinous ectoderm and lacks a mammalian-like mucus layer [91,92].

A possible source of host glycans for the insect hindgut microbiota is the peritrophic matrix, which acts as a barrier between the midgut absorptive tissues and the microbiota, analogous to a mammalian mucus layer [93]. In cockroaches, the peritrophic matrix is continually produced in the dorsal midgut and extruded along with digested food into the hindgut [91,92]. While peritrophic matrices can contain both O- and N-glycans [94], our data suggests a greater role of N-glycans in supporting the insect hindgut microbiota. We observed high expression of GH38 alpha-mannosidases, which have previously been shown to act on N-glycans [95,96]. Additionally, we identified other CAZymes associated with fucosidase, glucosidase, and galactosidase activity. Analysis of N-glycans in cockroach midgut preparations confirmed the presence of glycans containing these residues (Figure S10). These structures are also consistent with a published analysis of N-glycans in the peritrophic matrix of the Colorado potato beetle [94]. While expression of CAZymes active on O-glycans was much lower, *Bacteroides* did express genes to break down bonds present in O-glycan analyses of cockroach midguts (Figure S11). Overall, these results suggest that mannosylated N-glycans in the peritrophic matrix play a critical role in shaping the hindgut microbiota of cockroaches. Our hypothesis that the insect peritrophic matrix is a mechanism by which the host can shape the hindgut (as well as midgut) microbiota has implications for understanding hindgut microbial communities in other insects. As an example, honeybees represent another well-studied insect system with both a peritrophic matrix and a hindgut microbiota critical to host health [97–99]. An examination of data from the insect hindgut metagenome also suggests the presence of glycan-targeting CAZymes in the honeybee gut microbiota [100].

Finally, we note that for organisms further down the ‘microbial food web’, transcriptional activity remained remarkably stable across diets. While primary degraders exhibit large transcriptional shifts in response to changing diets, these efforts release a limited set of sugar monomers that, in turn, are processed via central metabolic pathways into a limited suite of fermentation end products. Despite having a comparable level of coverage in the metatranscriptome to that seen in some fiber degrading bacteria, sulfate reducing bacteria and methanogens, in particular, showed remarkable stability in transcriptional activity across diets. These organisms primarily survive on hydrogen and other fermentation products (short chain fatty acids and/or methylamines) released during the degradation of fiber by primary degraders. Our results suggest that the transcriptional plasticity exhibited by the fiber degraders is sufficient to maintain production of these end products at levels that support the continued persistence and activity of the rest of the gut microbiome, leading to the observed overall stability in gut microbiome composition seen overall.

While these results may seem contradictory to the many studies showing dietary impacts on gut microbiome composition, we would like to emphasize two points. First, we note that overall stability in gut microbiome seems to vary among organisms, with laboratory mice in particular showing strong responses to diet while the human gut microbiota frequently does not exhibit large structural changes that are greater than pre-existing inter-individual variability [28,31,32]. Further, many measures of microbial community composition and responses to dietary shifts are primarily aimed at identifying differences, while ignoring similarities. The goal of this study was to emphasize instead mechanisms that support gut microbiome homeostasis in the face of dietary differences.

## Conclusions

This work emphasized the importance of fiber degraders in supporting overall gut microbiome homeostasis. In our model, transcriptional plasticity in fiber degrading bacteria and – to a lesser extent– secondary fermenters was sufficient to maintain key hydrogenotrophic taxa at stable relative abundance and minimal alterations to their overall transcriptional profiles. Our results also suggest a role for host glycans present in the peritrophic matrix and passed into the hindgut as part of normal digestive function in providing a critical nutritional subsidy when the host diet lacks sufficient fiber sources. Together, these results provide new insight into host and microbial mechanisms supporting gut microbiome homeostasis in omnivorous cockroaches.

## Acknowledgements

This work was supported by the National Institute of General Medical Sciences of the National Institutes of Health (NIH) under award number R35GM133789. Glycan analyses were supported under the NIH award R24GH137782. We would like to thank Lauren Hudgens for help with the glycan analyses.

## Data Availability

Data associated with this study are available from NCBI SRA under BioProject accession number PRJNA994902. Scripts and workflow for the metatranscriptomic analyses can be found here: https://github.com/adepoy1/cockroach_metatranscriptome_manuscript.

