## Supplemental Text and Figures for "Metabolic flexibility among fiber degraders supports gut microbiome stability across host diets in the American cockroach"

**Contents**

1. Supplemental Methods: Glycan analysis 2
2. Supplemental Table Descriptions (Table S1-6, tables uploaded separately) 4
3. Supplemental Figures:
Supplemental Figure 1. Co-Occurrence Heatmaps for Lachnospiraeae 5
Supplemental Figure 2. Co-Occurrence Heatmaps for Oscillospiraceae 6
Supplemental Figure 3. Shannon diversity for metagenome and metatranscriptome 7
Supplemental Figure 4. Taxonomy and significantly enriched taxa for the metagenome and metatranscriptome datasets 8
Supplemental Figure 5. Family-level taxonomy bar plot for metagenome and Metatranscriptome 9
Supplemental Figure 6. Percent of Variation for RDA analyses 10
Supplemental Figure 7. Bar plot of total number of significantly changing genes for all taxa 11
Supplemental Figure 8. Relative abundance bar plot of main metabolic pathways 12
Supplemental Figure 9. Relative abundance of select chaperones and folding catalysts 13
Supplemental Figure 10. N-glycans from the cockroach midgut 14
Supplemental Figure 11. O-glycans from the cockroach midgut 14

#### 1 Supplemental Methods:

*Glycan analysis*

Midgut tissue from cockroaches fed microcrystalline cellulose (MCC) and a positive control sample of fetuin were used for glycan analyses. For N-glycan analysis, a 50 mM ammonium bicarbonate buffer was prepared, and the samples (in 900 µL buffer) were homogenized using a Bead Ruptor. The samples were transferred into a new tube and reduced with 5 mM DTT for 1 h at 55 °C. The samples were then carboxyamidomethylated with 15 mM iodoacetamide in the dark at room temperature for 45 min. The protein was washed with water and purified using a 10 kDa filtration membrane. The sample was resuspended in ammonium bicarbonate buffer and digestion with PNGaseF (37 ºC overnight) was performed to release N-glycans. Finally, the samples were then placed at 100 °C for 5 minutes to denature. The sample was cleaned using a preconditioned C18 Cartridge. The samples were then added to each of their respective columns, followed by the elution of the N-glycans of 3 mL of 5% acetic acid, which were then lyophilized for 45 minutes. The collected N-glycans were then permethylated (as with the linkage analysis) two times and spotted on the MALDI plate. The retained O-glycans were retained for further analysis.

For O-glycan linkage analysis, the same C18 cartridges described above (for N-glycan linkage analysis) were used. O-glycans were eluted and collected by adding 1 mL of 20%, 40%, 60%, and pure isopropyl alcohol (IPA) in 5% acetic acid in series. The collected O-glycans were then dried down. The O-glycopeptides from the samples, with fetuin being excluded, were beta eliminated by addition of a 19 mg/ml solution of sodium borohydride in 50mM NaOH overnight at 45 °C. The samples were then neutralized with the addition of 50 µL 10% acetic acid. This 50 µL addition of 10% acetic acid was repeated multiple times until the solution stopped reacting. Salt was removed via passage over an activated DOWX column ($H^+$). The samples were permethylated (as with the linkage analysis) and spotted on the MALDI plate. The O glycopeptides were also analyzed through LC-MS to confirm structures found in the O-glycans.

#### 2 Supplemental Table Descriptions

Table S1. Genomes used in constructing pangenomes for each group

Table S2. Metatranscriptome sequence quality control and annotation data

Table S3. Metagenome sequence quality control and annotation data

Table S4. DESeq2 results for taxa in Bacteroidota phylum

Table S5. DESeq2 results for taxa in Bacillota phylum

Table S6. DESeq2 results for taxa in Desulfobacterota and Euryarcheaota

### 3 Supplemental Figures:

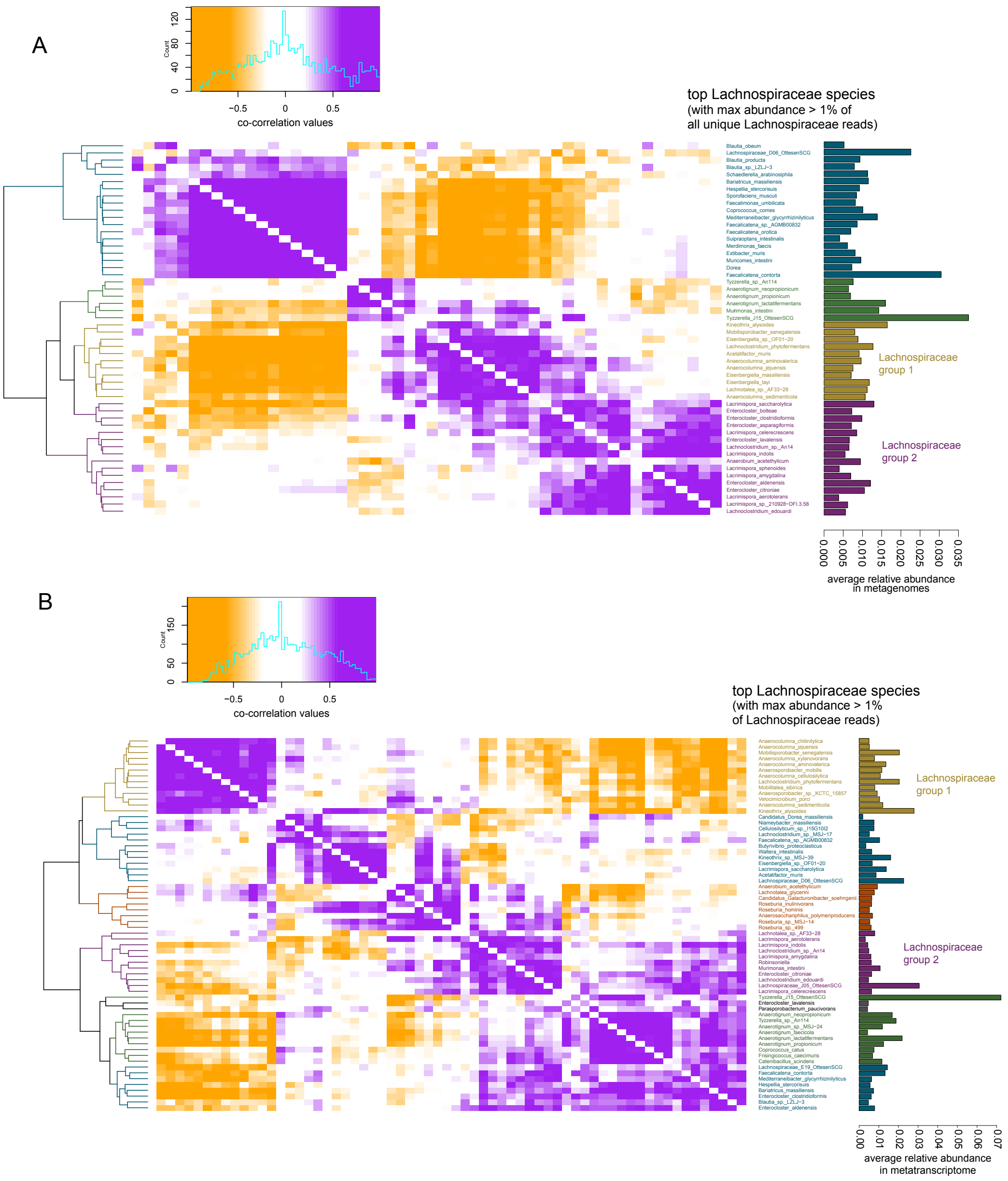


**Figure S1.** Co-occurrence heatmaps for genes from *Lachnospiraceae* in the metagenome (A) and metatranscriptome (B) dataset. Taxa displayed have a maximum relative abundance > 1% in any single sample.

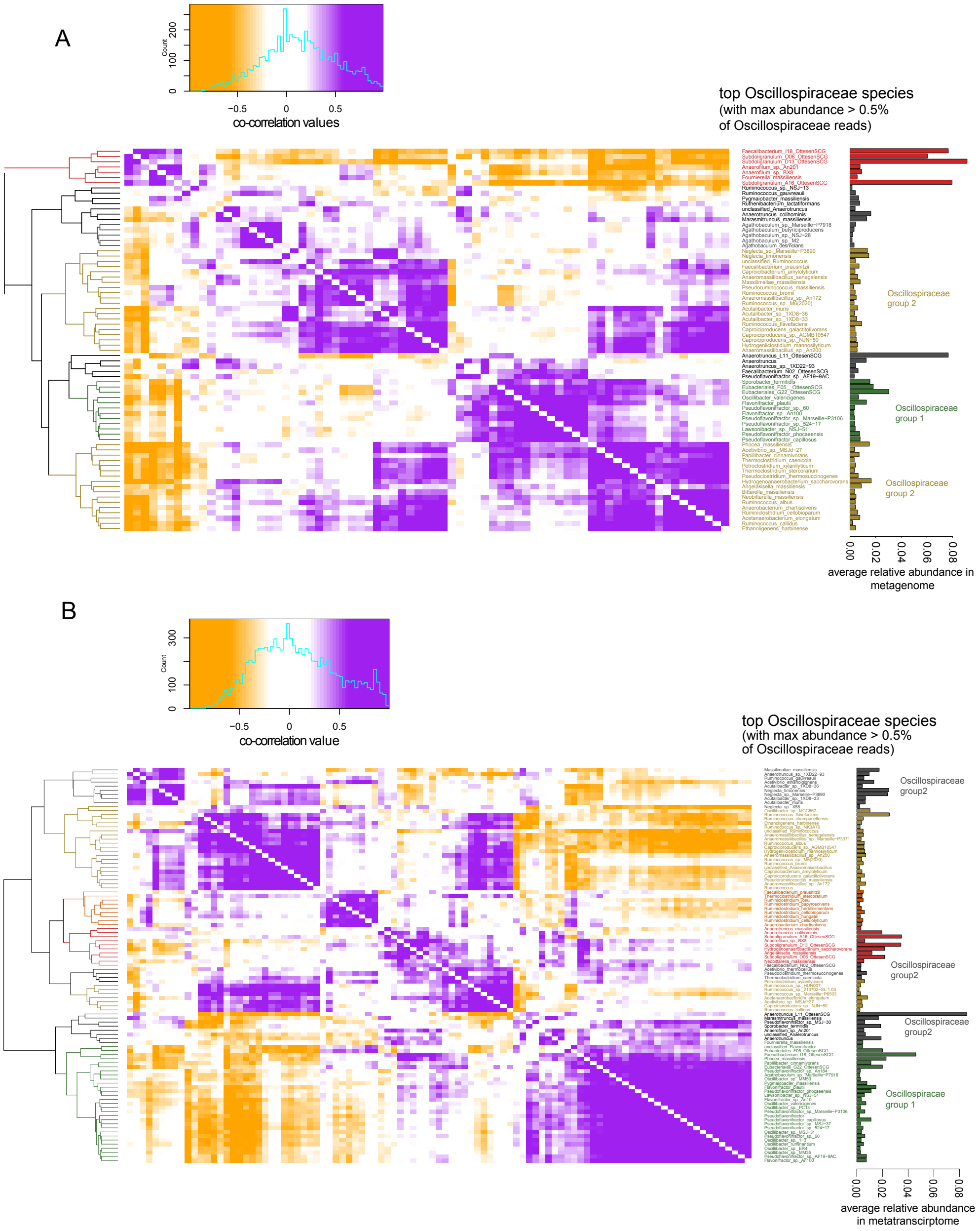


**Figure S2.** Co-occurrence heatmaps for genes from *Oscillospiraceae* in the metagenome (A) and metatranscriptome (B) dataset. Taxa displayed have a maximum relative abundance > 1% in any single sample.

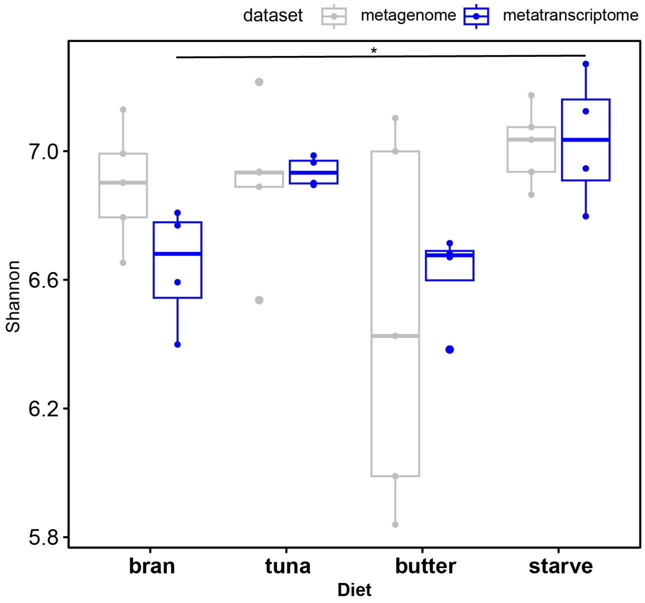


**Figure S3.** Shannon diversity metric for the metagenome (grey) and the metatranscriptome (blue) samples. Significance between diets was assessed using Kruskal-Wallis (for metagenome) or ANOVA (for metatranscriptome) and p-value cutoff of 0.05.

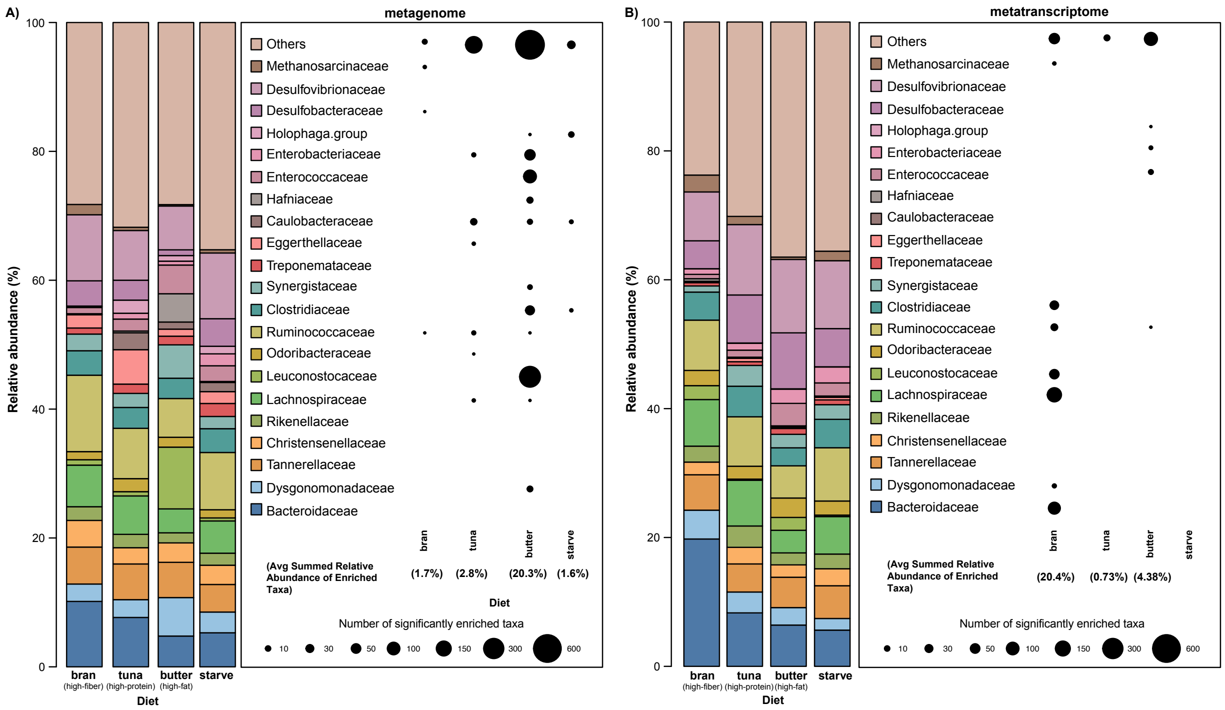


**Figure S4.** Taxonomy and significantly enriched taxa in each diet for the metagenome (A) and metatranscriptome (B) datasets. Relative abundance of families in the metagenome and metatranscriptome. Each column represents an average relative abundance of 5 replicates (metagenome) or 4 replicates (metatranscriptome) per diet. Families with a maximum relative abundance of at least 5% in any sample of either the metatranscriptome or metagenome dataset are included in the taxonomy. The number of significantly enriched taxa in each diet are enriched in each diet compared to all others (ex. bran vs. all other diets [tuna, butter, and starve], or high-fiber vs. low-fiber).

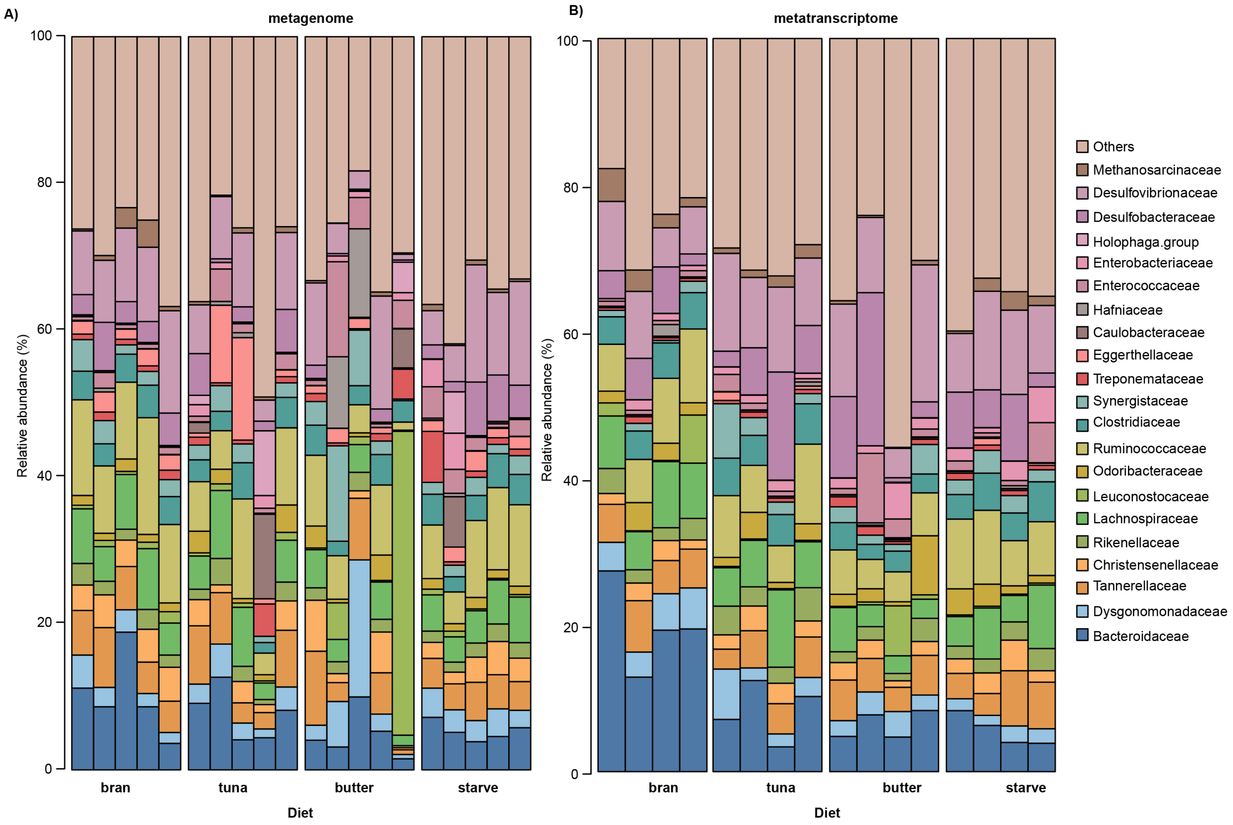


**Figure S5.** Relative abundance of families in the metagenome (A) and metatranscriptome (B). Each column represents a cockroach sample. Metagenomes have five replicates per diet and metatranscriptomes have four replicates per diet. Families with a maximum relative abundance of at least 5% in any sample of either the metatranscriptome or metagenome dataset are included in the taxonomy.

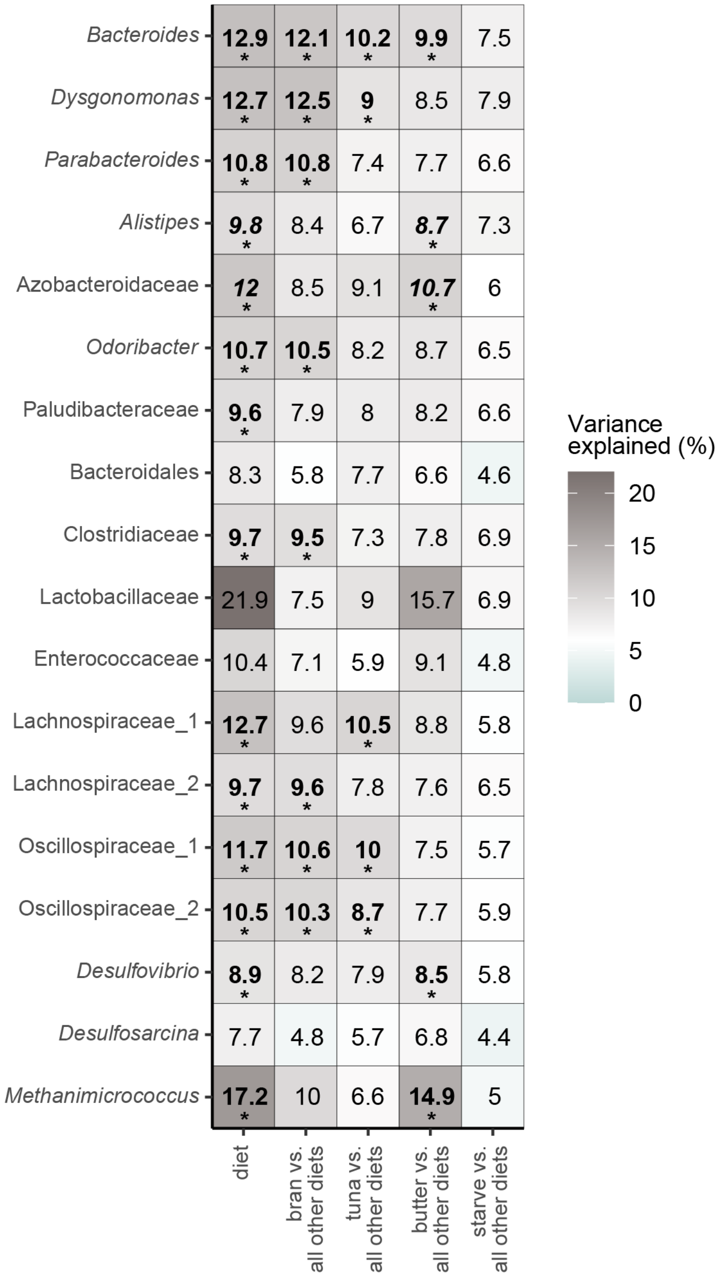


| | diet | bran vs. all other diets | tuna vs. all other diets | butter vs. all other diets | starve vs. all other diets |
|---|---|---|---|---|---|
| *Bacteroides* | **12.9*** | **12.1*** | **10.2*** | **9.9*** | 7.5 |
| *Dysgonomonas* | **12.7*** | **12.5*** | **9*** | 8.5 | 7.9 |
| *Parabacteroides* | **10.8*** | **10.8*** | 7.4 | 7.7 | 6.6 |
| *Alistipes* | ***9.8**** | 8.4 | 6.7 | ***8.7**** | 7.3 |
| Azobacteroidaceae | ***12**** | 8.5 | 9.1 | ***10.7**** | 6 |
| *Odoribacter* | **10.7*** | **10.5*** | 8.2 | 8.7 | 6.5 |
| Paludibacteraceae | **9.6*** | 7.9 | 8 | 8.2 | 6.6 |
| Bacteroidales | 8.3 | 5.8 | 7.7 | 6.6 | 4.6 |
| Clostridiaceae | **9.7*** | **9.5*** | 7.3 | 7.8 | 6.9 |
| Lactobacillaceae | 21.9 | 7.5 | 9 | 15.7 | 6.9 |
| Enterococcaceae | 10.4 | 7.1 | 5.9 | 9.1 | 4.8 |
| Lachnospiraceae_1 | **12.7*** | 9.6 | **10.5*** | 8.8 | 5.8 |
| Lachnospiraceae_2 | **9.7*** | **9.6*** | 7.8 | 7.6 | 6.5 |
| Oscillospiraceae_1 | **11.7*** | **10.6*** | **10*** | 7.5 | 5.7 |
| Oscillospiraceae_2 | **10.5*** | **10.3*** | **8.7*** | 7.7 | 5.9 |
| *Desulfovibrio* | **8.9*** | 8.2 | 7.9 | **8.5*** | 5.8 |
| *Desulfosarcina* | 7.7 | 4.8 | 5.7 | 6.8 | 4.4 |
| *Methanimicrococcus* | **17.2*** | 10 | 6.6 | **14.9*** | 5 |

**Figure S6.** Percent of variation explained using RDA for each taxa. Bold and starred (*) percentages represent significant using ANOVA and a significance cutoff of < 0.05. Data was normalized using a variance stabilized transformation using DESeq2.

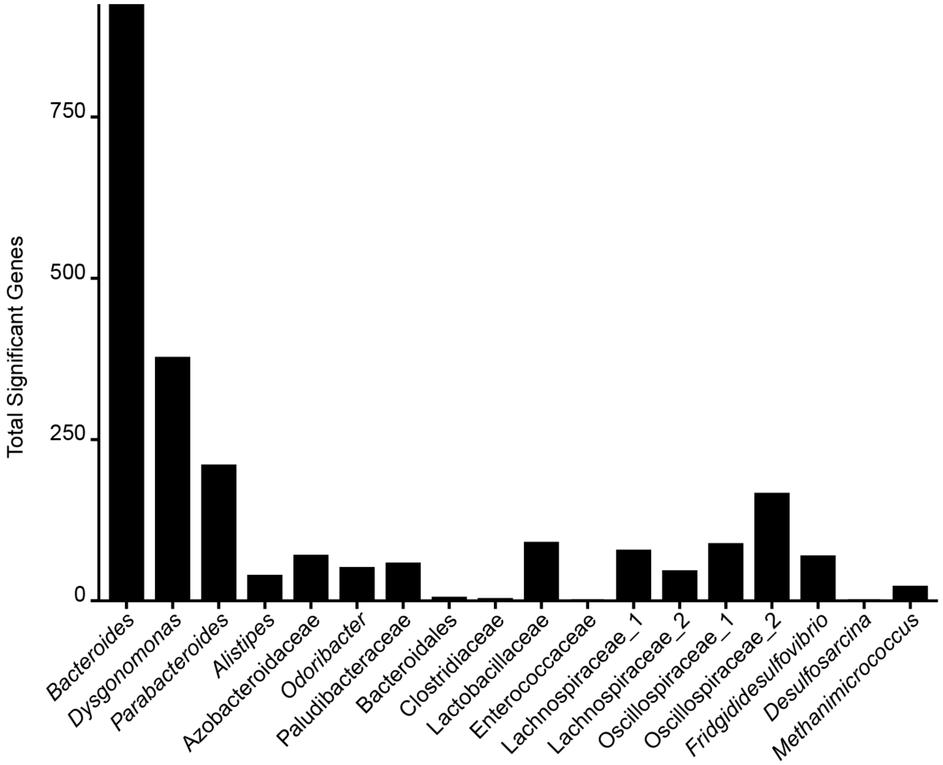


**Figure S7.** Bar plot of the total amount of significantly changing genes identified by DESeq2 for each taxon.

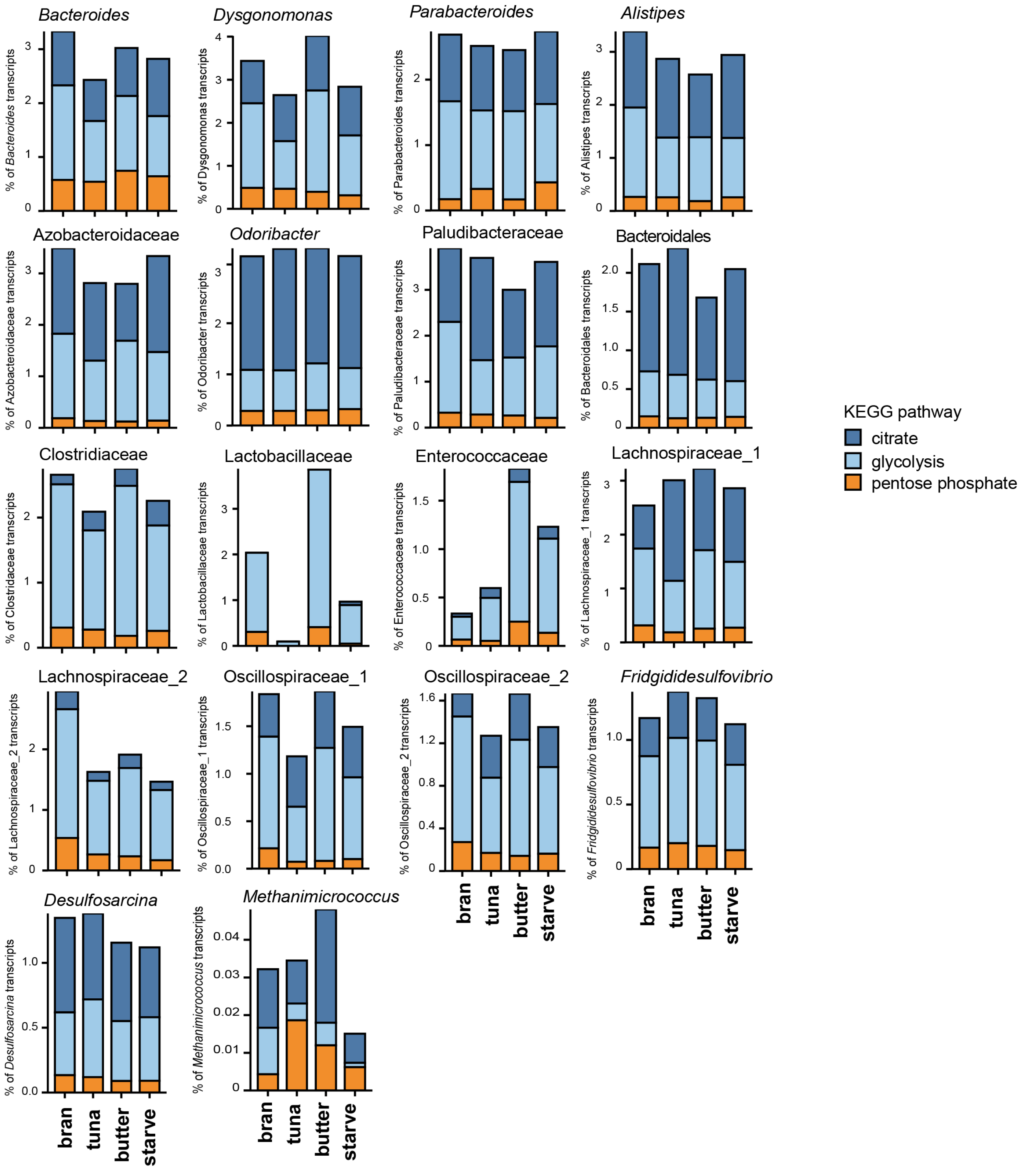


**Figure S8.** Average relative abundance of KEGG orthologs for major metabolic pathways (citrate, glycolysis, and pentose phosphate pathway). Relative abundances are expressed relative to each taxon's total number of reads per sample. Relative abundances were summed for each pathway before averaging. Averages were calculated for four replicates per diet.

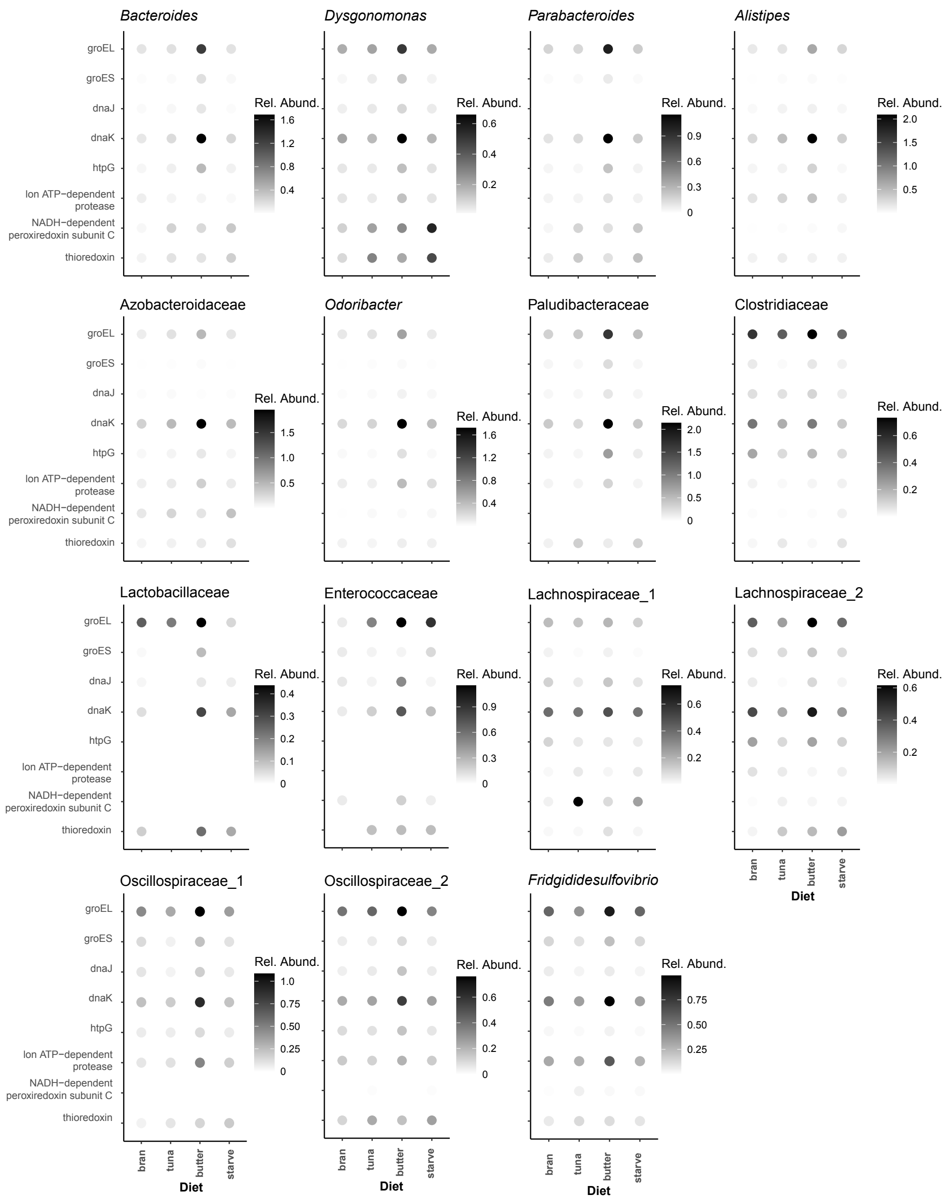


**Figure S9.** Relative abundance of select chaperones and folding catalysts. Relative Abundance is expressed relative to each taxon's total transcripts per sample. KO annotations for these genes were pulled based on high expression and DESeq2 results.

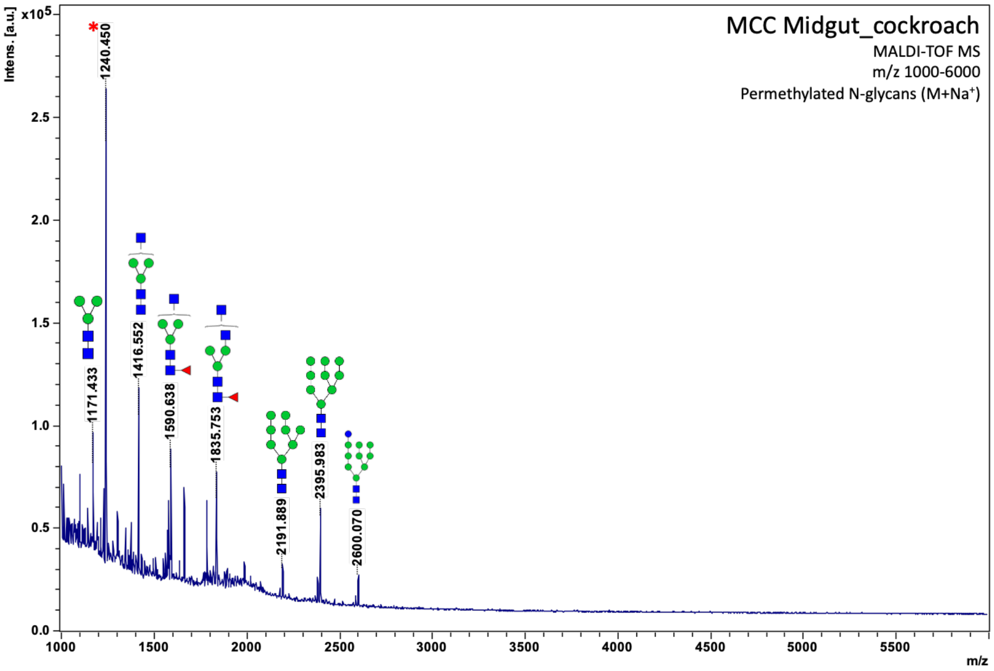


**Figure S10.** MALDI-TOF MS of permethylated N-glycans from cockroach midgut tissue.

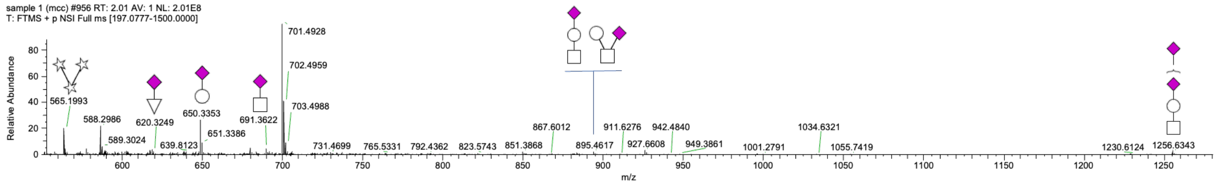


**Figure S11.** LC-MS of permethylated O-glycans from cockroach midgut tissue.
